# Single-nucleus transcriptomics of epicardial adipose tissue from females reveals exercise control of innate and adaptive immune cells

**DOI:** 10.1101/2023.11.02.565385

**Authors:** Irshad Ahmad, Shreyan Gupta, Patricia Faulkner, Destiny Mullens, Micah Thomas, Sharanee P. Sytha, Ivan Ivanov, James J. Cai, Cristine L. Heaps, Annie E. Newell-Fugate

**Author notes:** corresponding author: Annie E. Newell-Fugate **Email:**. Both authors have contributed equally (Irshad Ahmad, Shreyan Gupta). **Author Contributions:** Author contributions: C.L.H. and A.E.N.F research design; A.E.N.F., P.F., S.P.S., M.T., C.L.H. sample collection and preparation; I.A., S.G., data analysis; I.A., S.G., A.E.N.F., S.P.S., J.J.C., C.L.H. wrote the manuscript; I.A., S.G., A.E.N.F., S.P.S., J.J.C., D.M., I.I., C.L.H. edited the manuscript.

## Abstract

Coronary artery disease (CAD) is a leading cause of death in women. Although exercise mitigates CAD, the mechanisms by which exercise impacts epicardial adipose tissue (EAT) are unknown. We hypothesized that exercise promotes an anti-inflammatory microenvironment in EAT from female pigs. Yucatan pigs (n=7) were assigned to sedentary (Sed) or exercise (Ex) treatments and coronary arteries were occluded (O) with an ameroid to mimic CAD or remained non-occluded (N). EAT was collected for bulk and single nucleus transcriptomic sequencing (snRNA-seq). Exercise upregulated G-protein coupled receptor, S100 family, and FAK pathways and downregulated the coagulation pathway. Exercise increased the interaction between immune, endothelial, and mesenchymal cells in the insulin-like growth factor pathway and between endothelial and other cell types in the platelet endothelial cell adhesion molecule 1 pathway. Sub- clustering revealed nine cell types in EAT with fibroblast and macrophage populations predominant in O-Ex EAT and T cell population predominant in N-Ex EAT. Coronary occlusion impacted the largest number of genes in T and endothelial cells. Genes related to fatty acid metabolism were the most highly upregulated in non-immune cells from O-Ex EAT. Sub-clustering of endothelial cells revealed that N-Ex EAT separated from other treatments. In conclusion, aerobic exercise increased interaction amongst immune and mesenchymal and endothelial cells in female EAT. Exercise was minimally effective at reversing alterations in gene expression in endothelial and mesenchymal cells in EAT surrounding occluded arteries. These findings lay the foundation for future work focused on the impact of exercise on cell types in EAT.

**Significance Statement:** Coronary artery disease (CAD) is a leading cause of death in women. However, the role of epicardial adipose tissue (EAT) in the development of CAD in females and how exercise, which is recommended to slow CAD progression, impacts EAT are unknown. The effect of aerobic exercise on gene expression in EAT was investigated with RNA-sequencing, revealing significant alterations in fatty acid processing and immunoregulatory processes. This study provides valuable insights into the molecular and cellular changes induced in EAT by exercise in the context of chronic ischemic heart disease in females. These findings bolster current understanding of the impact of aerobic exercise on cardiac health in females and provide a foundation for future research in the field of exercise science.

## Introduction

Coronary artery disease (CAD) is the leading cause of death for women in the United States (U.S.) (1). Despite the large morbidity and mortality of CAD in American women, there is a general lack of awareness both of its impact on women’s health and the distinct sex-related disparities associated with this disease (2). Thus, there is a critical need to advance knowledge focused on the pathogenesis and management of CAD in women. In humans, 80% of the heart surface area is covered by epicardial adipose tissue (EAT) which secretes cytokines to modulate physiological and pathophysiological processes in the coronary arteries and myocardium (3, 4). Moreover, EAT is involved in the breakdown of fatty acids, serving as a local energy source for cardiomyocytes during periods of increased energy demand (5). Despite EAT’s beneficial functions, increased EAT is detrimental to both the structure and function of heart (6). Altered secretion of adipokines and proinflammatory molecules from EAT causes increased instability of atherosclerotic plaques (7–10).Thickened EAT also is associated with atrial enlargement, impaired diastolic filling, elevated myocardial lipid content and lipotoxic injury to cardiomyocytes, and cardiac remodeling (6). Furthermore, myocardial infiltration with epicardial adipocytes and the release of cytokines into the myocardial layer results in tissue inflammation and cardiac muscle dysfunction (11). Thus, imbalance between the protective and harmful effects of EAT contributes to CAD progression.

Increased physical activity is recommended for the prevention and secondary treatment of CAD. Aerobic exercise mitigates CAD via enhanced cardiac function and vascular adaptations which increase tissue blood flow to meet metabolic demands (12). Furthermore, aerobic exercise reduces EAT volume in females (13–15). Yet, how aerobic exercise modulates molecular mechanisms in EAT is unknown. In this study, we dissected the opposing effects of exercise and sedentary lifestyle on EAT at both the tissue and single-cell level in female Yucatan miniature swine with chronic ischemic heart disease (**Fig. 1 A, B, D**). The impact of aerobic exercise on gene expression in EAT was investigated with bulk RNA-sequencing (RNA-seq), revealing significant alterations in fatty acid processing and immunoregulatory processes. Single-nuclei transcriptome (snRNA-seq) analysis of EAT further unveiled diverse molecular signatures for both non-immune and immune cells, along with tissue-level gene and pathway modifications due to exercise. This study highlighted the influence of aerobic exercise on intercellular communication pathways, such as insulin-like growth factor (IGF) and platelet endothelial cell adhesion molecule 1 (PECAM1). Furthermore, differential gene expression (DGE) analysis revealed that genes related to cell adhesion, migration, fatty acid metabolism, and angiogenesis were modulated in non-immune and immune cells, depending on exercise status. This study provides valuable insights into the molecular and cellular changes induced in EAT by aerobic exercise in the context of ischemic heart disease in females.

**Figure 1.**
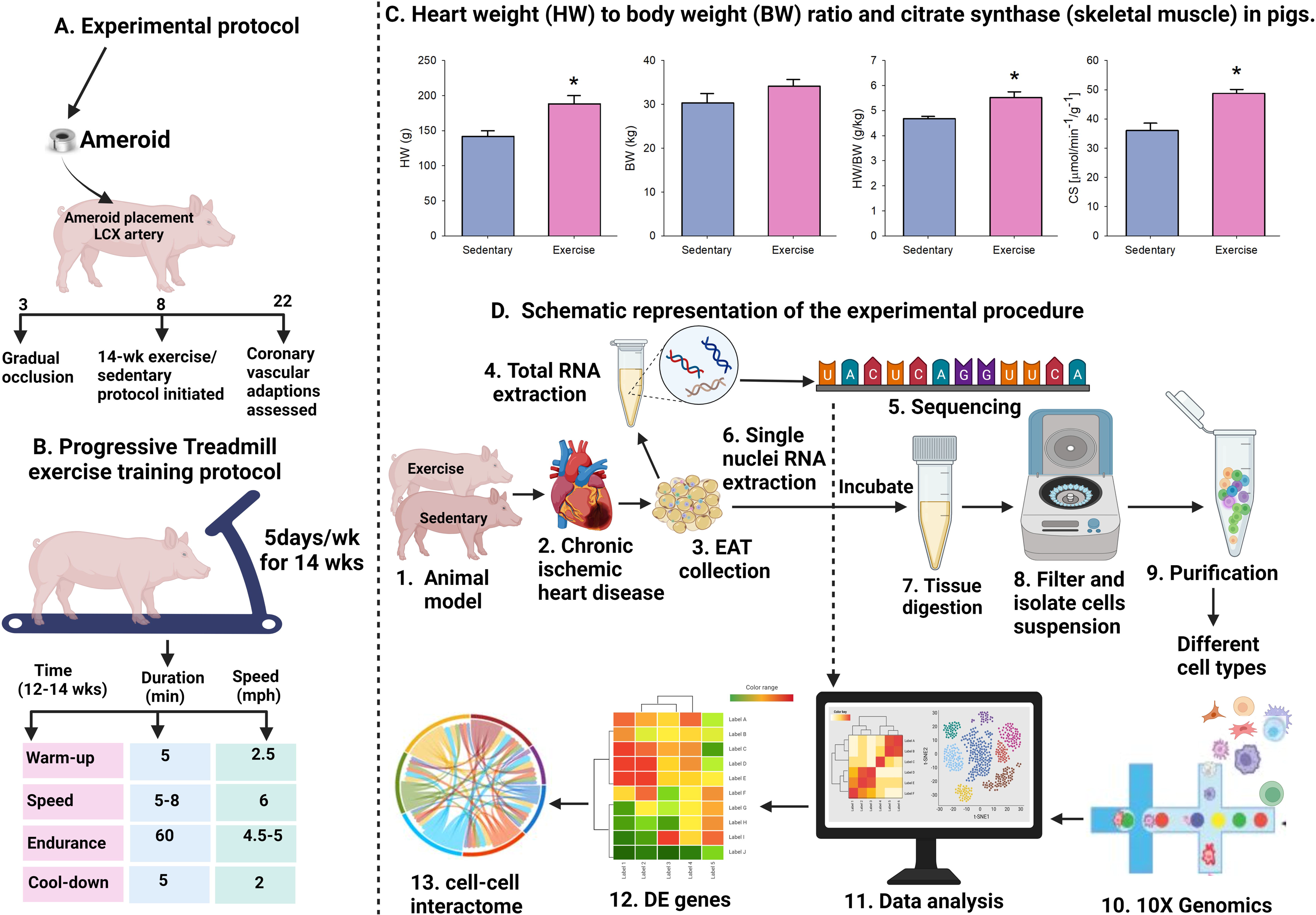
Schematic representation of the experimental protocol followed during the experiment. (A) Overall animal study design (B) exercise training regimen (C) Heart to body weight ratio and citrate synthase in female pigs at the end of the experiment (D) Bulk and single nuclei RNA extraction and computational analysis. This image was created using BioRender. CS: citrate synthase; HW: heart weight; BW: body weight; LCX: left circumflex coronary artery; EAT: epicardial adipose tissue; DE: differentially expressed. n=4 per treatment group; * *p* < 0.05.

## Results

### Fourteen-week aerobic exercise training regimen increases heart to body weight ratio and skeletal muscle citrate synthase activity in female pigs

Effectiveness of the 14-wk aerobic exercise-training program was assessed by comparison of heart-to-body weight ratio and skeletal muscle oxidative enzyme activity in exercise-trained and sedentary female pigs (**Fig. 1C**). Although body weight did not differ between exercise-trained and sedentary pigs, the heart-to-body weight ratio was greater in exercise-trained pigs. Citrate synthase activity was greater in the medial and lateral heads of the triceps brachii muscle in exercise-trained as opposed to sedentary pigs (**Fig. 1C)**.

### Aerobic exercise upregulates pathways and processes related to cellular migration, proliferation, differentiation, and transduction in EAT

We performed RNA-seq on EAT from exercise-trained and sedentary female pigs. Canonical pathways upregulated in order of most to least significance were: G- protein coupled receptor (GPCR) signaling, S100 family, focal adhesion kinase (FAK) signaling, cAMP response element binding protein (CREB), Fcy receptor mediated phagocytosis, phagosome formation, and interleukin 13 (IL-13) (**Fig. 2A**). The pathways with the strongest positive z-score indicating the strongest upregulation in response to exercise were: S100 family, CREB, Fcy receptor mediated phagocytosis, and phagosome formation. Pathways downregulated in response to exercise in order of most to least significance were: coagulation system, cyclic AMP (cAMP)-mediated signaling, G alpha subunits 12/13, and interleukin 4 (IL-4). The pathway with the strongest negative z-score indicating the strongest downregulation in response to exercise was: coagulation system (**Fig. 2A**). A heat map of the diseases and processes affected by aerobic exercise demonstrated that cellular movement followed by immune cell trafficking were the most upregulated processes in response to exercise (**Fig. 2B**). On the other hand, lipid metabolism, hematological system development, inflammatory response, and cell-to-cell signaling have genes associated with them that were both up- and down-regulated in response to exercise (**Fig. 2B**).

**Figure 2.**
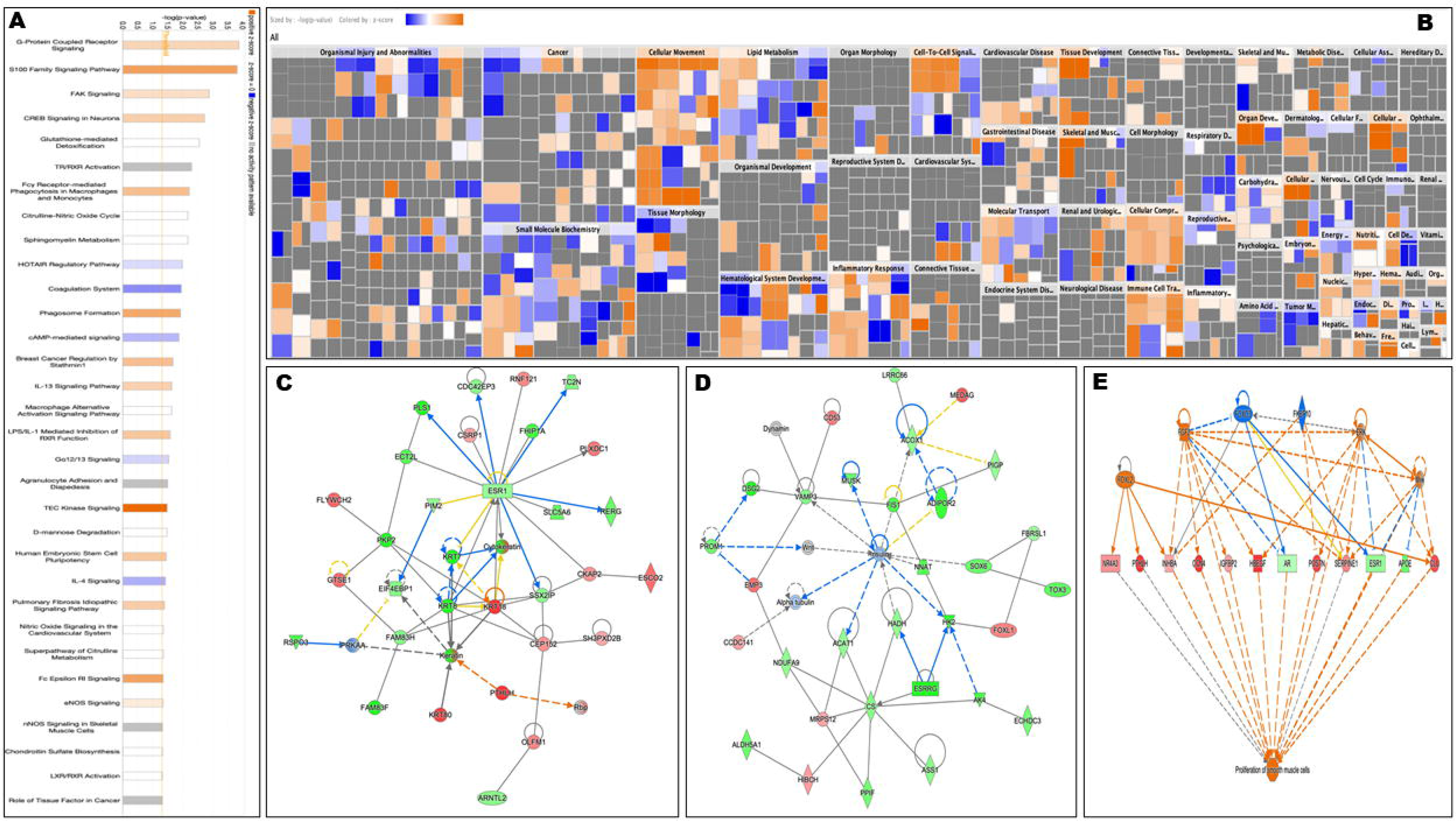
Pathways and diseases in epicardial adipose tissue of female pigs affected by exercise. (A) Top 30 pathways found in bulk adipose tissue to be significantly affected by exercise in female pigs. Orange: predicted upregulation; Blue: predicted downregulation; White: no effect. (B) Heat map of diseases and processes in bulk adipose tissue to be significantly affected by exercise training in female pigs. Orange: increased effect; Blue: decreased effect; White: no effect. (C) Estrogen receptor alpha network in bulk adipose tissue impacted by exercise. (D) Insulin signaling network in bulk adipose tissue impacted by exercise. (E) Predicted regulators of smooth muscle cell proliferation in bulk adipose tissue affected by exercise. Green: decreased gene expression; Red: increased gene expression; Blue: predicted gene inhibition; Orange: predicted gene activation

Several of the top pathways with multiple genes up- and/or down-regulated in their network analysis were: estrogen receptor alpha (ESR1) (**Fig. 2C**), insulin signaling and fatty acid metabolism (**Fig. 2D**), and smooth muscle cell proliferation (**Fig. 2E**). *ESR1* was downregulated and signaling downstream of *ESR1* included both down- (solute carrier family 5 member 6 (*SLC5A6*), cytokeratin, keratin 7 (*KRT7*), pim-2 proto- oncogene (*PIM2*), plastin 1 (*PLS1*), epithelial cell transforming 2 like (*ECT2L*), FHF complex subunit HOOK interacting protein 1A (*FHIP1A*) and up-regulated (cysteine and glycine rich protein 1 (*CSRP1*), plexin domain containing 1 (PDXC1), cytoskeleton associated protein 2 (*CKAP2*), keratin 18 (*KRT18*)) genes (**Fig. 2C**). Genes in the insulin signaling and fatty acid metabolism pathway were predominantly related to fatty acid oxidation and mitochondrial function and were downregulated in response to exercise: muscle associated receptor tyrosine kinase (*MUSK*), mitochondrial fission 1 (*FIS1*), adiponectin receptor 2 (*ADIPOR2*), acyl-coA oxidase 1 (*ACOX1*), vesicle associated membrane protein 3 (*VAMP3*), neuronatin (*NNAT*), acetyl-coA acetyl transferase 1 (*ACAT1*), hydroxyacyl-coA dehydrogenase (*HADH*), estrogen related receptor gamma (*ESRRG*) (**Fig. 2D**). Genes in the smooth muscle cell proliferation pathway were predominantly upregulated in response to exercise - (insulin-like growth factor binding protein 2 (*IGFBP2*), serpine family E number 1 (*SERPINE1*), cellular communication network factor 4 (*CCN4*), periostin (*POSTN*), heparin binding EGF-like growth factor (*HBEGF*)) - with the exception of androgen receptor (*AR*), *ESR1,* and apolipoprotein E (*APOE*) which were downregulated (**Fig. 2E**).

### Aerobic exercise increases interaction amongst immune and mesenchymal cells in the IGF pathway and between endothelial cells, fibroblasts, and other cell types in adhesion molecule pathways

We isolated single-nuclei suspensions from non-occluded (N) and occluded (O) regions of EAT from exercise-trained (Ex) and sedentary (Sed) swine, resulting in four experimental groups: N-Ex, O-Ex, N-Sed, and O-Sed. Single-nuclei suspensions were profiled using 10x Genomics Chromium droplet snRNA-seq. The resulting EAT single nuclei atlas had 24,382 individual nuclei that were visualized with uniform manifold approximation and projection (UMAP) and grouped into 17 clusters. We performed unsupervised clustering on the cells using the Louvain algorithm in the Seurat R package (16). The clustering analysis revealed nine distinct clusters (**SI Appendix Fig. 1**). These clusters were annotated using markers from the pig atlas (https://dreamapp.biomed.au.dk/pigatlas/) with specific markers linked to each cellular phenotype to identify cell types. Finally, nine unique cell types were identified based on differential expression of marker genes in the nuclei clusters (**Fig. 3A**, **SI Appendix Dataset S1**). Cells identified included: 1) beige adipocytes (receptor accessory protein 1*(REEP1),* prospero homeobox 1 *(PROX1),* multimerin 1(*MMRN1*), semaphorin 6A (*SEMA6A*), par-6 family cell polarity regulator gamma (*PARD6G*); 2) endothelial cells (*PECAM1),* fms related receptor tyrosine kinase 1*(FLT1),* adhesion G protein-coupled receptor *(L4ADGRL4),* cysteine and tyrosine rich 1 *(CYYR1),* dedicator of cytokinesis 9

**Figure 3.**
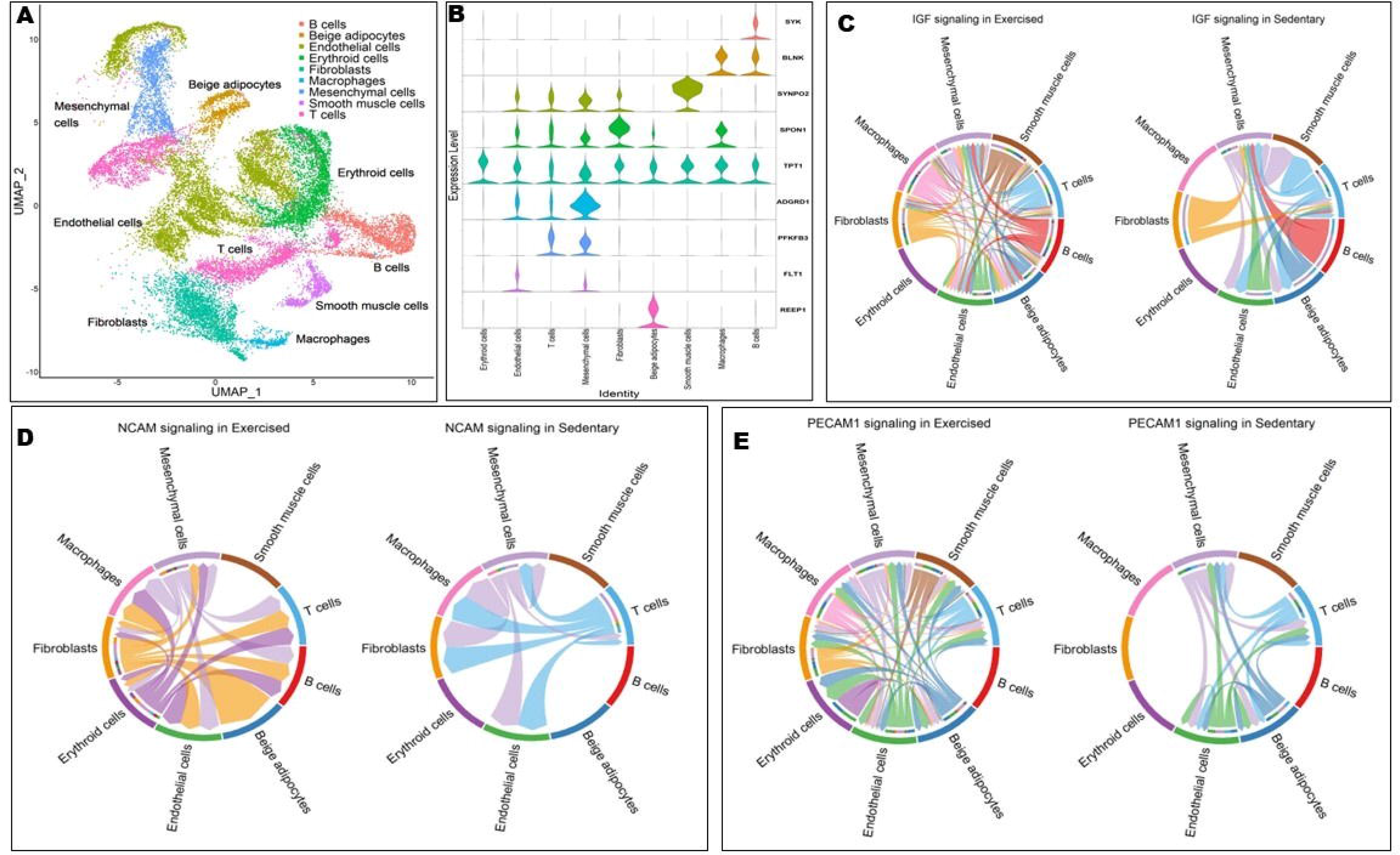
Single nuclei gene expression from epicardial adipose tissue of sedentary and exercised female pigs. (A) Single nuclei clustering in epicardial adipose tissue from exercise-trained and sedentary female pigs. (B) Expression of established marker genes for each cell type in each cluster. Circle plots representing most prominent cell-cell communications between exercise and sedentary groups in the insulin-like growth factor (IGF) (C), neural cell adhesion molecule (NCAM) (D), and platelet endothelial cell adhesion molecule 1 (PECAM1) (E) signaling pathways.

*(DOCK9),* scavenger receptor class A member 5 *(SCARA5),* zinc finger and BTB domain containing 7C *(ZBTB7C),* phospholipase D1 *(PLD1),* protein tyrosine phosphatase receptor type B *(PTPRB),* CD93 molecule *(CD93),* apolipoprotein A1 *(APOA1),* regulator of G protein signaling 5 *(RGS5),* receptor activity modifying protein 2 *(RAMP2),* glycosylphosphatidylinositol anchored high density lipoprotein binding protein 1 *(GPIHBP1),* LIM domain binding 2 *(LDB2*); 3) fibroblasts (spondin 1 (*SPON1),* gelsolin *(GSN*); 4) mesenchymal cells (adhesion G protein-coupled receptor D1 (*ADGRD1*); 5) smooth muscle cells (synaptopodin 2 (*SYNPO2*); 6) erythroid cells (tumor protein, translationally-controlled 1 (*TPT1*); 7) macrophages (colony stimulating factor 1 receptor (*CSF1R),* B cell linke*r (BLNK),* amyloid beta precursor protein binding family B member 1 interacting protein *(APBB1IP),* CD163 molecule *(CD163),* B cell scaffold protein with ankyrin repeats 1 *(BANK1),* stabilin 1 *(STAB1*); 8) B cells (spleen associated tyrosine kinase *(SYK),* vav guanine nucleotide exchange factor 3 *(VAV3*), and 9) T cells (6- phosphofructo-2-kinase/fructose-2,6-biphosphatase 3 *(PFKFB3),* IKAROS family zinc finger 3 *(IKZF3),* protein tyrosine phosphatase receptor type c *(PTPRC),* Src kinase associated phosphoprotein 1 *(SKAP1),* talin 2 *(TLN2*) (**Fig. 3B**, **SI Appendix Table S1**).

Intercellular communication between non-immune cells (endothelial cells, beige adipocytes, fibroblast, mesenchymal cells, smooth muscles cells and erythroid) and immune cells (macrophages, B cells, T cells) was analyzed via CellChat algorithm (17). Aerobic exercise impacted the following cell-to-cell communication signaling pathways: IGF, neural cell adhesion molecule (NCAM), and PECAM1. For IGF signaling, all immune cell classes had the largest number of interactions with endothelial and mesenchymal cells in Ex-EAT. In contrast, only B and T cells heavily communicated with mesenchymal cells in Sed-EAT (**Fig. 3C**). For NCAM, a glycoprotein in the immunoglobulin superfamily, the strongest communication strength was observed between fibroblast and erythroid cells with all other cell types in Ex-EAT from exercise trained pigs. T cells and mesenchymal cells had the greatest number of communications with other cell types in Sed-EAT (**Fig. 3D**). PECAM1 is expressed at endothelial cell intercellular junctions and regulates leukocyte trafficking. In Ex-EAT, endothelial cells had increased interactions between all cell types except for B cells. In Sed-EAT, only T cells, endothelial cells, mesenchymal cells, and beige adipocytes interacted (**Fig. 3E**).

### Aerobic exercise upregulates cellular adhesion and metabolism genes in endothelial cells, adipocytes, smooth muscle cells, fibroblasts, and B cells

DGE analysis was performed using EdgeR between Ex-EAT and Sed-EAT. The top 10 differentially expressed genes for each cell type were sorted based on average log fold change. The DGE for each cell type are shown in **SI Appendix Dataset S2** and **SI Appendix Fig. 2**. For non-immune cells, DEGs included ELOVL fatty acid elongase 6 (*ELOVL6),* acetyl-CoA carboxylase alpha *(ACACA),* malic enzyme 1 *(ME1),* acyl-CoA *(ACYL),* stearoyl-CoA desaturase *(SCD).* Immune cells in Sed-EAT had differentially expressed genes related to immunoregulation and basic cellular processes (i.e. FKBP prolyl isomerase 5 (*FKBP5),* glutamate-ammonia ligase *(GLUL))*. *ELOVL6* was upregulated across all non-immune cell types except for mesenchymal cells in Ex- EAT. All immune cell categories in Sed-EAT had increased gene expression of *FKBP5* which is involved in protein folding and trafficking.

### Comprehensive profiling of cell type-specific gene expression reveals distinct molecular signatures in EAT in response to exercise and coronary artery occlusion status

When experimental groups were divided into N-Sed, O-Sed, N-Ex, and O-Ex, all nine cell type clusters were found (**Fig. 4A**). O-Ex tissue had abundant fibroblasts while N-Ex tissue had a large amount of T cells and endothelial cells (**Fig. 4B**). O-Sed tissue had abundant endothelial cells while N-Sed tissue had many endothelial and erythroid cells (**Fig. 4B**). The fibroblast and macrophage cell populations primarily were comprised of cells from the O-Ex treatment. On the other hand, the B cell, adipocyte, erythroid cell, and mesenchymal cell populations were predominantly comprised of cells from the N- Sed treatment. Most of the T cell cluster was from the N-Ex treatment (**Fig. 4B and SI Appendix Table S2**).

**Figure 4.**
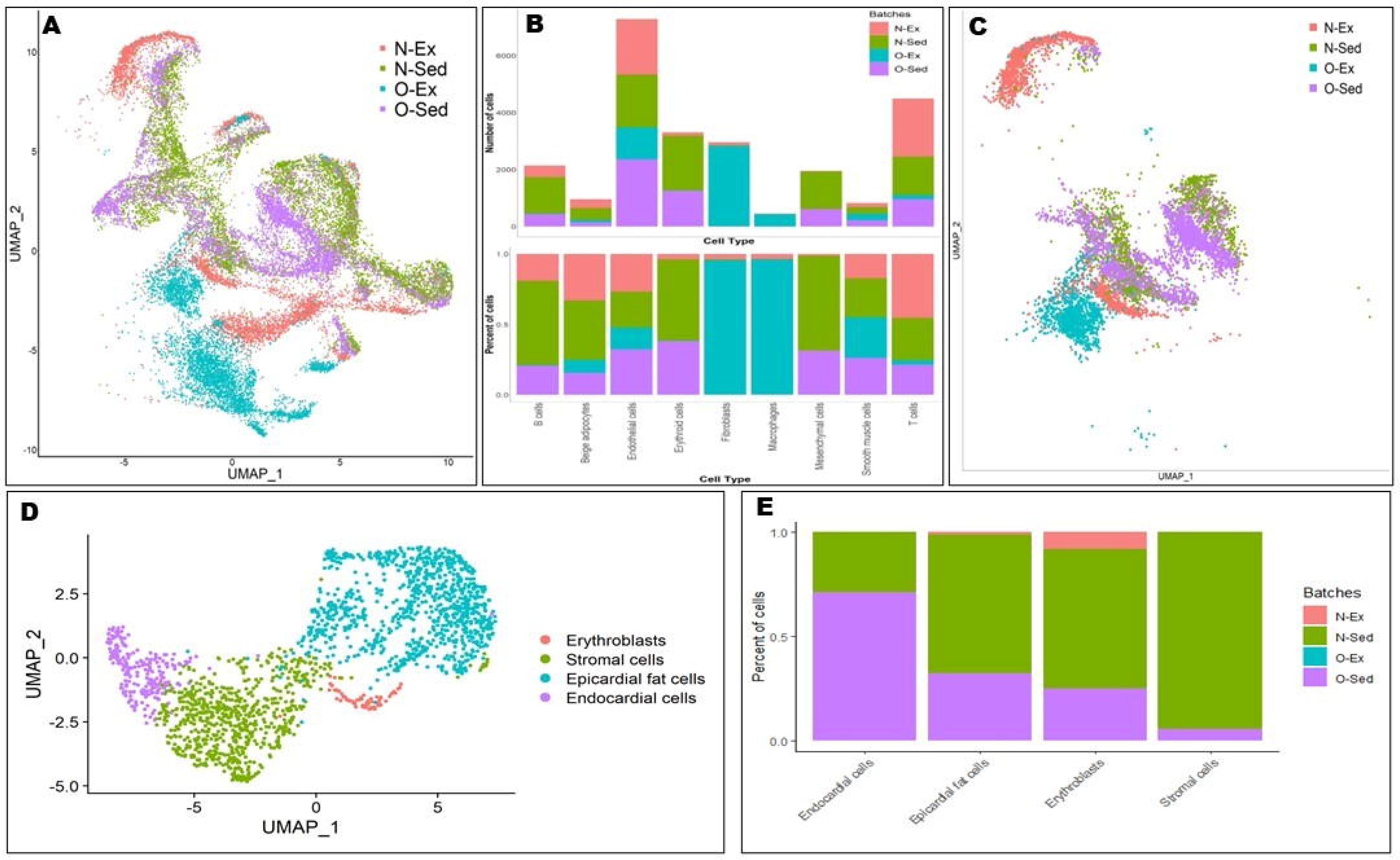
Single nuclei gene expression from epicardial adipose tissue surrounding occluded and non-occluded coronary arteries from sedentary and exercise trained female pigs. (A) Occluded and non-occluded unbiased clustering of 24,382 cells revealed nine different cell clusters. (B) Number and percentage of cell types in each treatment (exercise trained and sedentary) and batch-wise distribution in each occlusion*exercise treatment (occluded and non-occluded) expressed genes. (C) Subclustering of endothelial cells showed separation of N-Ex group from all other treatments. (D) Sub-clustering of mesenchymal cells showed four different cell types. (E) Percentage of each cell type in the mesenchymal subcluster by occlusion*exercise treatment. N: non-occluded; O: occluded; Ex: exercise trained; Sed: sedentary.

Samples from N-Sed and O-Sed found genes involved in cell adhesion, migration processes, and fatty acid metabolism were modulated in both non-immune and immune cells. All cells except for fibroblasts and macrophages in N-Sed EAT had upregulated expression of fibronectin 1 (*FN1*), which is involved in cell adhesion and migration processes (**SI Appendix Dataset S3, SI Appendix Fig. 3**). On the other hand, fibroblasts in N-Sed EAT showed the highest levels of *ME1,* IQ motif containing GTPase activating protein (*IQGAP2),* and phospholipid scramblase (*PLSCR)* which are involved in fatty and lipid movement and metabolism. Macrophages in N-Sed EAT had a unique gene expression profile with increased gene expression of only FAT atypical cadherin (*FAT3)* and glutamate ionotropic receptor kainate type 4 (*GRIK4*). O-Sed EAT had highly variable results for DGE among different cell types (**SI Appendix Dataset S3, SI Appendix Fig. 3**). Non-immune cells from O-Sed EAT had upregulated expression of leucine-rich repeat-containing protein (*LRRC43)*, while B cells from O-Sed EAT had upregulated expression of *IQGAP2,* colony stimulating factor 1 receptor (*CSF1R*), limbic system associated membrane protein (*LSAMP*), and stabilin 1 (*STAB1*) (**SI Appendix Dataset S3, SI Appendix Fig. 3**). T cells had upregulation of 3-oxoacid coA-transferase 1 (*OXCT1*), talin 2 (*TLN2*), pyruvate dehydrogenase kinase 4 (*PDK4*), sorbin and SH3 domain containing 1 (*SORBS1*), growth hormone receptor (*GHR*), diacylglycerol O- acyltransferase 2 (*DGAT2*), phosphodiesterase 3B (*PDE3B*), and *ELOLV6* (**SI Appendix Dataset S3, SI Appendix Fig. 3**).

Genes involved in fatty acid metabolism (*ELOLV6, DGAT2*, lipoprotein lipase (*LPL*), fatty acid binding protein (*AFABP*)) and angiogenesis (matrix gla protein (*MGP*), *GHR*, colony stimulating factor 1 (*CSF1*), microtubule actin crosslinking factor 1 (*MCAF1*)) were upregulated in non-immune cells in O-Ex EAT (**SI Appendix Dataset S4, SI Appendix Fig. 4**). Mesenchymal cells in O-Ex EAT had the most robust upregulation of these genes. Endothelial cells in N-Ex EAT had many upregulated genes such as phospholipase C episilon 1 (*PLCE1*), semaphorin 3C (*SEMA3C*), basonuclin zinc finger protein 2 (*BNC2*), phospholipase D 1 (*PLD1*), cyclin dependent kinase 14 (*CDK14*), glycoprotein M6A (*GPM6A*) and cell adhesion associated, oncogene regulated (*CDON*). The percentage of non-immune cells in EAT expressing genes related to metabolism (i.e. *ELOVL6 and* acetyl-coA carboxylase alpha *(ACACA*)) was higher in O- Ex compared to N-Ex pigs (**SI Appendix Dataset S4, SI Appendix Fig. 4**). In N-Ex EAT, macrophages had upregulation of *CDON,* EBF transcription factor 1 *(EBF1),* DLC1 Rho GTPase activating protein *(DLC1)*. In contrast, T and B cells in O-Ex EAT had upregulation of genes similar to non-immune cells in O-Ex EAT (**SI Appendix Dataset S4, SI Appendix Fig. 4)**. In T cells, 9,786 genes in non-occluded EAT and 2,037 genes in occluded EAT were upregulated **(SI Appendix Fig. 5)**. In endothelial cells, 3,179 in non-occluded EAT and 2,045 genes in occluded EAT were upregulated (**SI Appendix Fig. 5)**.

### Aerobic exercise does not reverse alterations in gene expression in EAT endothelial cells due to coronary artery occlusion and minimally impacts EAT mesenchymal cell gene expression

Sub-clustering of endothelial cells resolved by the four treatment groups resulted in five clusters with separation of the N-Ex group from the rest of the treatment groups (**Fig. 4C**). Furthermore, N-Ex group had upregulated gene expression of latent transforming growth factor beta binding protein 2 (*LTBP2*)*, PKHD1 Like 1 (PKHD1L1), FERM Domain (FRMD), CDON,* and glycoprotein M6A *(GPM6A),* which are all related to cell adhesion and structure (**SI Appendix Fig. 6 and SI Appendix Dataset S5**). Upon analysis of the mesenchymal cell cluster based on differential expression of marker genes, four different cell types were identified: endocardial, epicardial fat cells, erythroblasts, and stromal cells (**Fig. 4D**). Of these identified cell types, most mesenchymal cells were epicardial fat cells. When the subclustered mesenchymal cells were analyzed by treatment group, the only cells associated with N-Ex were epicardial fat cells and erythroblasts (**Fig. 4E**). The stromal cell subcluster was almost completely comprised of cells from N-Sed. Most endocardial cells were attributed to the O-Sed group. Surprisingly, O-Ex contributed very little to the mesenchymal cell population (**Fig. 4E**).

## Discussion

This study examined the cellular composition and gene expression of EAT in female pigs in response to aerobic exercise and experimental ischemic heart disease. We applied both RNA-seq and snRNA-seq to gain novel insights into the diversity of cell types in EAT and their responses to exercise and coronary artery occlusion. RNA-seq revealed that aerobic exercise upregulated pathways related to cell movement, signal transduction, and proliferation and processes related to macrophage function such as phagosome formation and phagocytosis. These findings do not completely correspond with those in rodents and humans, which have found that exercise decreases inflammation (18), particularly due to macrophages (19), and increases angiogenesis (20, 21) in subcutaneous (sc) white adipose tissue (WAT). However, most of the studies on the impact of exercise on scWAT have been done in obese subjects and many rodent studies only included males. Interestingly, aerobic exercise caused both up- and down- regulation of genes related to lipid metabolism and the inflammatory response, particularly with respect to the interaction of T cells and macrophages in EAT. We applied snRNA-seq to gain a nuanced understanding of the impact of both aerobic exercise and coronary artery occlusion on cell types and gene expression in female EAT.

A sedentary lifestyle caused accumulation of B cells and mesenchymal cells in EAT. In obesity, B cells accumulate in WAT and interact with T cells to produce proinflammatory cytokines (22). Moreover, in scWAT of obese patients macrophage populations shift from anti-inflammatory M2 cells to pro-inflammatory M1 cells (23). Surprisingly, short term aerobic exercise decreases M2 macrophage levels in scWAT and visceral (v) WAT of obese male mice (24). In contrast to what is found in WAT of obese male mice in response to acute exercise, macrophage and T cell numbers were greater in Ex-EAT from female pigs. These results suggest that aerobic exercise impacts inflammation in EAT from females quite differently from scWAT or perhaps highlights the differences between WAT in mice and pigs.

As aerobic exercise upregulated expression of genes involved in fatty acid synthesis in B cells, but not in T cells or in macrophages, B cells may play a larger role in immunometabolism in EAT than these other two immune cells. This finding is in direct contrast to scWAT where macrophages are one of the main immune cell populations in control of immunometabolism (25). Mesenchymal stem cells from both humans and mice stimulate regulatory B cell functions via cell-to-cell contact, soluble factors, and extracellular vesicles which lead to anti-inflammatory reactions (26). In obesity, adipose- derived mesenchymal cells exhibit pro-inflammatory properties, attract inflammatory immune cells, and create an inflammatory microenvironment which causes mesenchymal cell dysfunction (27). Additionally, B cells accumulate in obese WAT and interact with T cells to produce proinflammatory cytokines (22). An increased cell-to-cell interaction in the IGF1 pathway between mesenchymal cells, B cells, and T cells in Sed- EAT indicates that a sedentary lifestyle is proinflammatory. In Sed-EAT, T cells interact with mesenchymal cells in the PECAM1 and NCAM pathways which indicates T cells are critical to cell movement and adhesion in Sed-EAT. These findings suggest that adaptive immune cells may serve as a dominant communication “hub” in Sed-EAT and that immune cell function in EAT is quite different from that in other WAT depots.

Coronary artery occlusion interacted with exercise status in EAT to modulate cell number and gene expression. Surprisingly, coronary artery occlusion increased fibrosis and macrophage infiltration in Ex-EAT as opposed to Sed-EAT. Chronic aerobic exercise in males decreases infiltration of M1 macrophages and favors recruitment of M2 macrophages in scWAT (28). It is possible that coronary artery occlusion counteracts the beneficial effects of chronic exercise on WAT inflammation. It is also possible that the effects of chronic exercise on EAT are different from the effects on other WAT depots. With respect to occlusion status and exercise, N-Ex EAT had large numbers of T cells and endothelial cells. It is not surprising that exercise stimulated increased numbers of endothelial cells in N-Ex EAT, because exercise induces WAT angiogenesis (29). Our study is the first report of the effects of chronic exercise training on T cell numbers in WAT from any depot. Isolation and immunotyping of T cells from EAT would be the next step to determine the mechanistic implications of this finding. Contrary to what was expected, both N-Sed and O-Sed EAT had upregulation of genes related to cell adhesion and migration. These findings suggest that coronary artery occlusion has minimal effect on these processes in Sed-EAT. However, coronary artery occlusion does result in upregulation of genes related to fatty acid metabolism in T cells from Sed-EAT, which again suggests a critical role for T cells in this EAT.

*ELOVL6*, which is involved in the elongation of long-chain fatty acids (30), was upregulated across all non-immune cell types except mesenchymal cells in Ex-EAT. Interestingly, *ELOVL6* was upregulated in these same cell types as well as B and T cells in O-Ex EAT. *ACACA* and *ACLY,* which are involved in fatty acid synthesis, also were upregulated across all non-immune cell types and B cells in Ex-EAT. However, occlusion status did not affect the expression of these two genes. These findings suggest that exercise may increase the production of fatty acids in most cell types in female EAT. Increased expression of these genes also occurs in scWAT from lean and Roux-en-Y gastric bypass (RYGB) obese patients, which indicates a preference for lipogenesis as opposed to lipolysis in these states (31). Other studies have found exercise decreases lipogenesis in scWAT of obese female rodents (32) and increases lipolysis in the scWAT of obese and lean men (33). Increased expression of *FKBP5*, which is a co-chaperone that modulates glucocorticoid action (34), in all immune cell classes in Sed-EAT indicates that the immune system is highly controlled by cortisol in sedentary states. In animal studies, deletion or inhibition of *FKBP5* causes decreased WAT mass and protection against diet-induced weight gain, insulin resistance, and hepatic steatosis (35, 36). In humans, *FKBP5* expression in abdominal scWAT correlates positively with markers of insulin resistance and type 2 diabetes (37). Therefore, glucocorticoids seem to have similar effects in all WAT depots, including EAT.

With respect to the interaction of occlusion and exercise, in non-immune cells, especially mesenchymal cells, from O-Ex EAT genes associated with fatty acid metabolism and angiogenesis were upregulated. In contrast, in N-Ex EAT endothelial cells had upregulation of genes involved in cell adhesion and movement. One of the most striking findings in this study was the large number of upregulated genes in T cells in N-EAT, implicating T cells as important to the normal function of EAT in females. There was a five-fold decrease in the number of upregulated genes in T cells in EAT upon coronary artery occlusion. These findings suggest that T cells may be critical to the normal function of EAT in females. Another remarkable finding was the separation of endothelial cells in N-Ex EAT from endothelial cells in EAT from all other treatments. Given that the gene signature of endothelial cells in O-Ex EAT is similar to the gene signatures of O-Sed and N-Sed EAT suggests that coronary artery occlusion blocks the potential beneficial effects of aerobic exercise on endothelial cells in EAT. Endothelial cells are essential for vascular function, the delivery of nutrients and oxygen, and removal of wastes from EAT (38). Therefore, this finding coupled with the fact that O-Ex EAT has more fibrosis and macrophage infiltration indicates that coronary artery occlusion in females is extremely detrimental to the health of the surrounding EAT. Moreover, chronic aerobic exercise is unable to counteract these deleterious effects. Sub-clustering of mesenchymal cells found that most of the cells in this population were from Sed-EAT. In high fat diet fed mice, mesenchymal stem cells can differentiate into adipocytes but exercise reverses this effect (39). Our findings with respect to mesenchymal cells correspond with study and support the ability of exercise to limit differentiation of the EAT mesenchymal cell pool into epicardial fat cells.

In conclusion, this study highlights that chronic aerobic exercise, independent of coronary artery occlusion status, increases interaction between immune and mesenchymal and endothelial cells in EAT from female pigs. Furthermore, genes related to cell adhesion and metabolism in these cell types as well as adipocytes and smooth muscle cells are upregulated in response to exercise. T cells appear to be critical to the normal function of female EAT, whereas B cells are the immune cell class in female EAT most sensitive to aerobic exercise. Surprisingly, chronic aerobic exercise is minimally effective at reversing alterations in gene expression in endothelial and mesenchymal cells in EAT surrounding occluded coronary arteries in female pigs. These findings lay the foundation for future work focused on the mechanistic impact of exercise on various cell types in EAT from women.

## Materials and Methods

### Experimental animals and surgical instrumentation

Animal protocols were approved by the Texas A&M University Institutional Animal Care and Use Committee and conformed to the National Institutes of Health (NIH) “Guide for Care and Use of Laboratory Animals, 8^th^ edition,” revised 2011). Adult Yucatan miniature swine (6-7 months of age) were surgically instrumented with an ameroid constrictor around the proximal left circumflex coronary artery (**SI Appendix**).

### Sedentary and Exercise Protocols

Exercise trained pigs underwent a progressive treadmill exercise training program, 5 days/week for 14 weeks, as described previously (40) (**SI Appendix**).

### Isolation of coronary epicardial adipose tissue

Following completion of the 14-week exercise training or sedentary protocol, pigs were humanely euthanized and epicardial perivascular fat was sectioned from both the non-occluded left anterior descending coronary artery and the collateral-dependent left circumflex coronary artery. Perivascular fat was snap-frozen in liquid nitrogen and subsequently stored at -80 °C for later analysis. Subsequent visual inspection of the ameroid occluder during dissection of the left circumflex artery under a dissection microscope during dissection of the left circumflex artery indicated 100% occlusion in all pigs used in this study.

### Bulk transcriptomics

EAT (occluded and non-occluded tissue pooled) from exercise-trained (n=4) and sedentary (n=3) female pigs had total RNA extracted with TRIzol® (Thermofisher Scientific, Waltham, MA). Total RNA was quantified (Nanodrop 3300, Thermofisher, Wilmington, DE) followed by bioanalysis (Agilent 2100 Bioanalyzer, Agilent Technologies, Inc., Santa Clara, CA) for RNA quantity and quality. RNA integrity scores fell within the following range: 6.2–9.1. cDNA libraries were prepared, and sequencing was performed at the Molecular Genomics Core, Texas A&M University.

### Single-nucleus RNA sequencing

Isolation of EAT nuclei was performed following the “Daughter of Frankenstein protocol for nuclei isolation from fresh and frozen tissues using OptiPrep continuous gradient V.2” (41). Nuclei were resuspended in 0.1% BSA in PBS and immediately processed for the generation of single-nuclei RNA libraries using the microdroplet-based RNA. Nuclei samples were diluted if necessary to a target concentration of between 500 – 1,500 nuclei/µL and used for single nuclei RNA sequencing library preparation (**SI Appendix**).

### Bioinformatic Analysis

UMI count matrices for each single-cell sample were generated by aligning reads to the genome (Landrace_pig_v1 (GCA_001700215.1) using 10X Genomics Cell Ranger software. The DGE analysis was performed using edgeR package (42). Genes that did not occur frequent enough were filtered out, with a cutoff of at least 100 counts per million. Significantly differentially expressed genes were determined using a P-value cutoff of 0.01. The functional enrichment of top 100 key differentially expressed genes (DEGs) was carried out using PANTHER.db (https://www.pantherdb.org/) to identify key enriched pathways. For within-and cross-tissue communication prediction, UMI count matrices and cell type/state assignment were exported for each cell as two input files for CellChat R package (**SI Appendix**).

## Acknowledgments

We would like to thank Jeff Bray and Andrew Hillhouse for their assistance with this work.

## Data, Materials and Software Availability

Bulk RNA-seq and snRNA-seq data are deposited into the BioSample database under accession number <u>GSE246709</u>. Supplementary Data for the present findings are available within Supporting Information files. Scripts for data processing and downstream analyses are available through GitHub at https://github.com/Xenon8778/Pig_EAT_scRNAseq

---

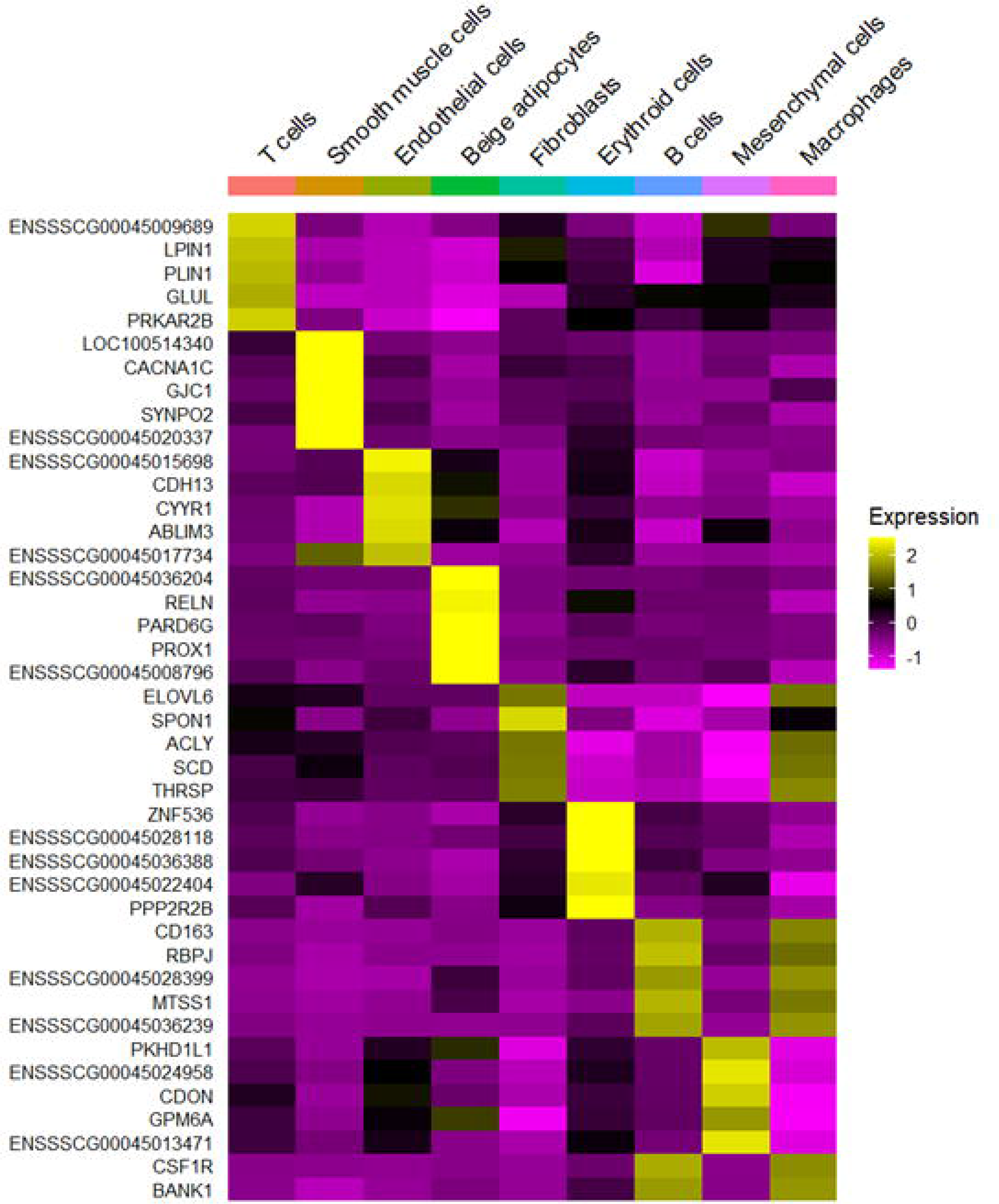

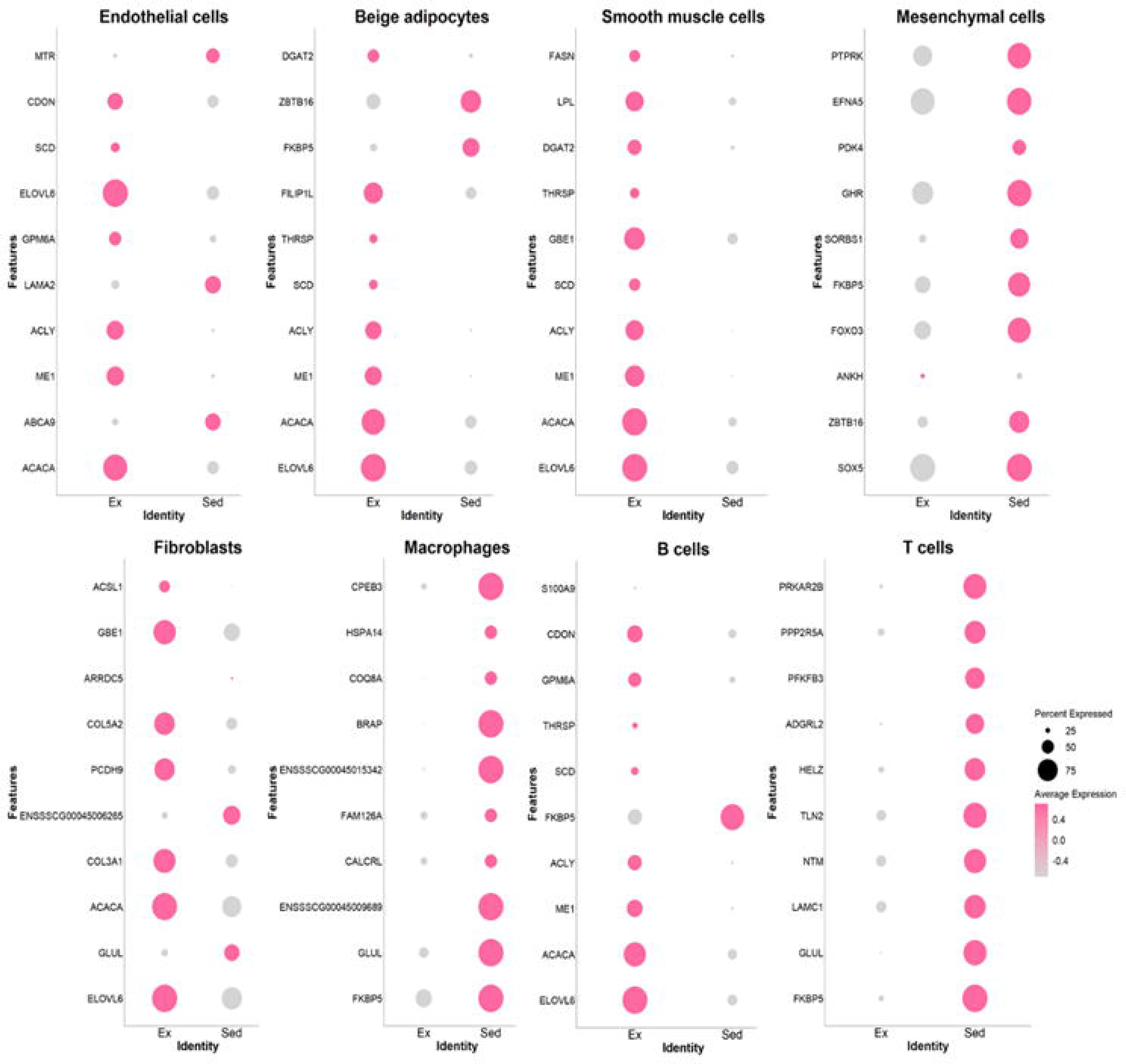

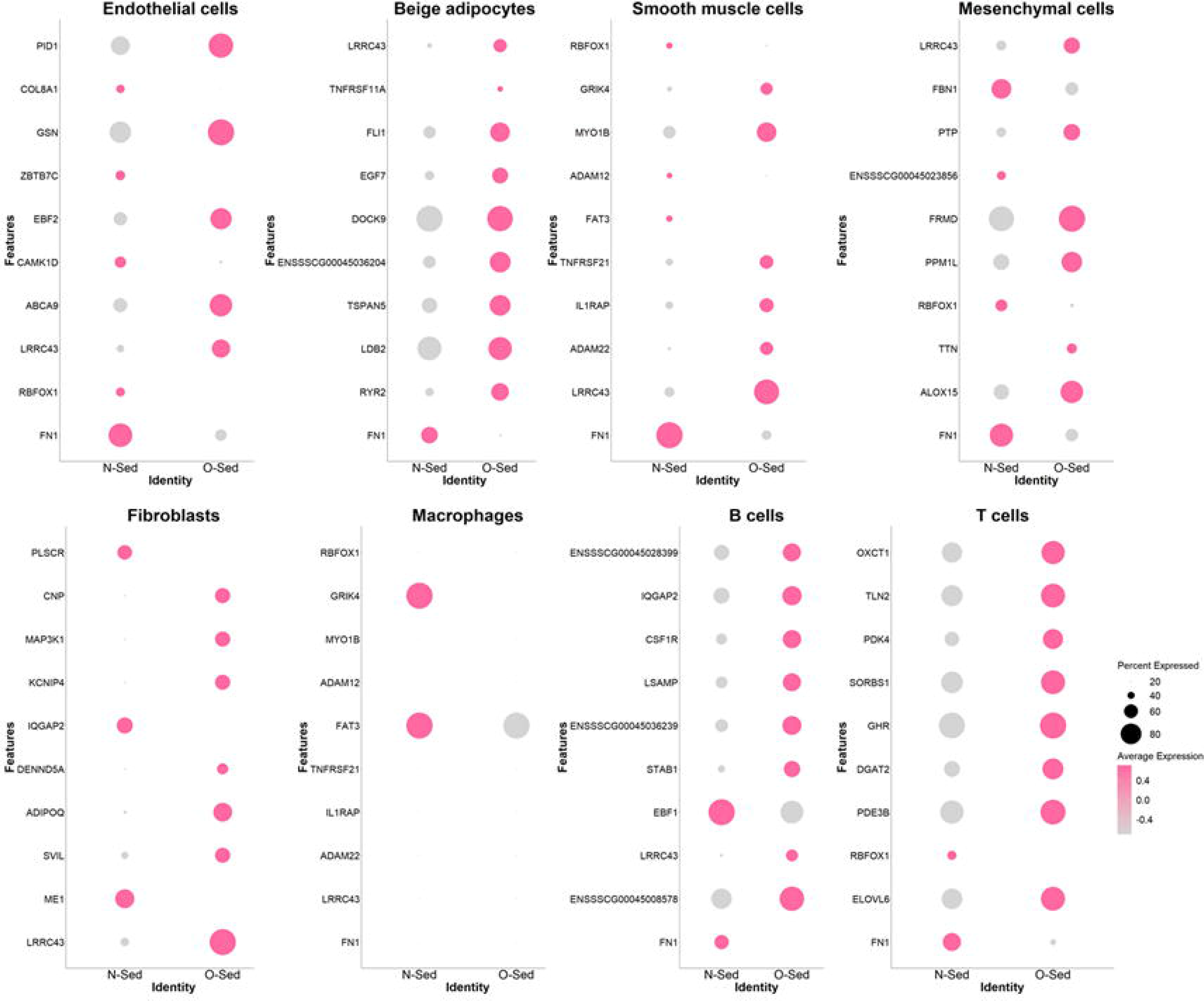

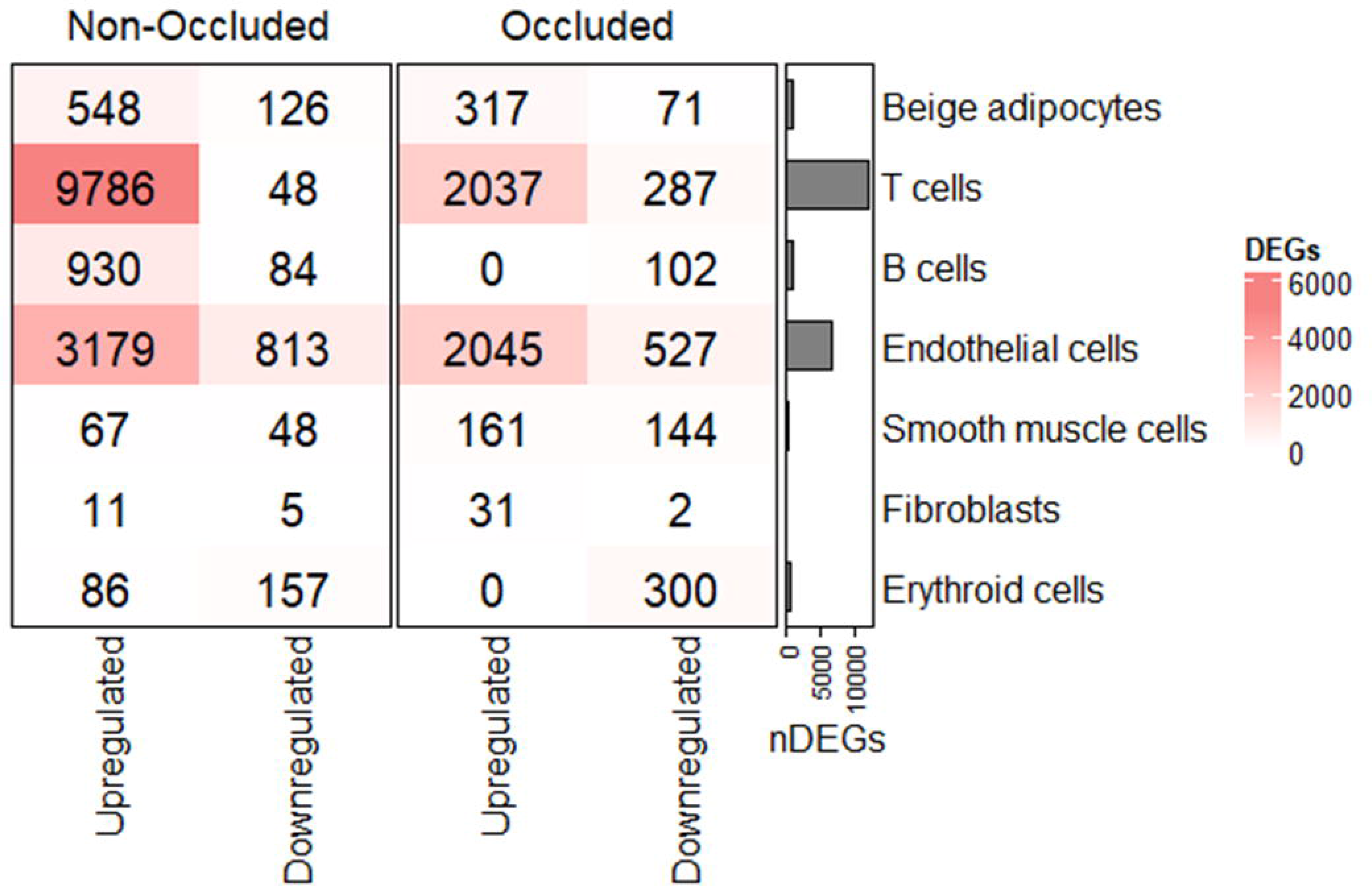

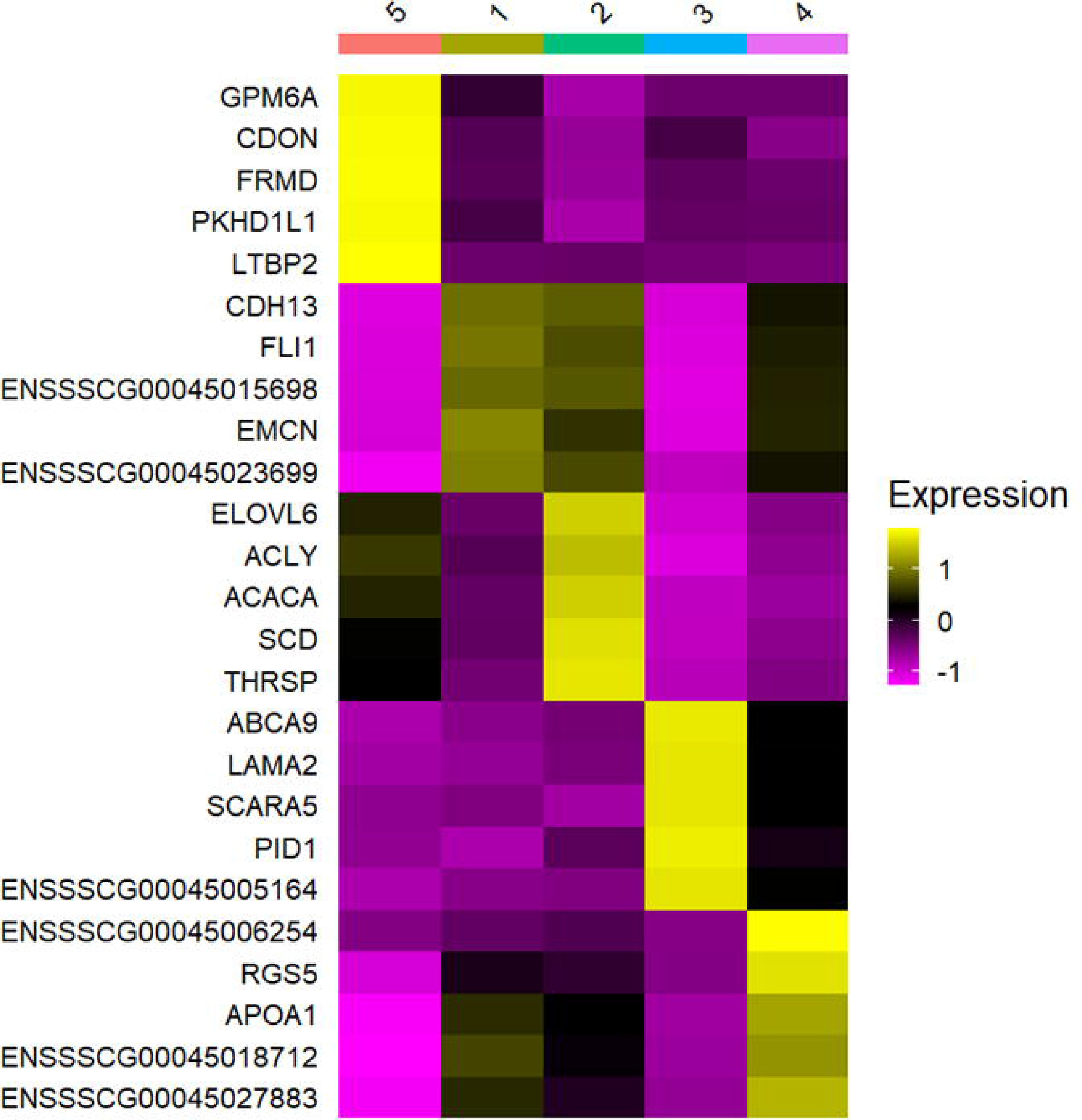

