## Supplemental Figure Legends for "Single-nucleus transcriptomics of epicardial adipose tissue from females reveals exercise control of innate and adaptive immune cells"

**Supplementary Figure Legends**

**Supplementary Figure S1: Expression of top transcript markers for each cell type found in single cell sequencing of epicardial adipose tissue from exercise-trained and sedentary female pigs.** Yellow is upregulated and magenta is downregulated expression level of the top 42 genes expressed in all cell types. As this study was conducted in porcine epicardial adipose tissue, many of the genes have not been identified by homologous name and function to human genes and, as such, are shown by their Ensembl Stable Id. Refer to supplementary dataset S1.

**Supplementary Figure S2.** **Dot plots of differential gene expression within each cell type in epicardial adipose tissue from exercised and sedentary female pigs**. Pink represents upregulated and gray represents downregulated transcript expression level. The size of the dot represents the percentage of cells expressing the gene in each cell type in exercise-trained and sedentary groups. Ex: exercise-trained; Sed: sedentary. Refer to supplementary dataset 2.

**Supplementary Figure S3: Dot plots of the differential gene expression within each cell type in epicardial adipose tissue surrounding occluded or non-occluded coronary arteries from sedentary female pigs.** Pink represents upregulated and gray represents downregulated transcript expression level. The size of the dot represents the percentage of cells expressing the gene in each cell type in sedentary non-occluded and occluded groups. N-Sed: non-occluded sedentary. O-Sed: occluded sedentary. Refer to supplementary dataset 3.

**Supplementary Figure S4**: **Dot plots of the differential gene expression within each cell type in epicardial adipose tissue surrounding occluded or non-occluded coronary arteries from exercise-trained female pigs.** Pink represents upregulated and gray represents downregulated transcript expression level. The size of the dot represents the percentage of cells expressing the gene in each cell type in exercised non-occluded and occluded groups. N-Ex: non-occluded exercise-trained. O-Ex: occluded exercise-trained. Refer to supplementary dataset 4.

**Supplementary Figure S5. Total number of upregulated and down regulated genes in epicardial adipose tissue surrounding occluded and non-occluded coronary arteries independent of exercise status.** Intensity of the red color corresponds to numbers of differentially expressed genes (DEGs) with red being the largest number DEG (6000) and light pink to white being the smallest number of DGE (0). The length of the gray bar next to each cell type corresponds to the total number of upregulated or downregulated DEGs by cell type. The largest numbers of genes were altered in T cells and endothelial cell.

**Supplementary Figure S6: Expression of top transcript markers specifically for endothelial cells in different subclusters found in single cell sequencing of epicardial adipose tissue from exercise-trained and sedentary female pigs.** Yellow is upregulated and magenta is downregulated expression level of the top 25 genes expressed in all cell types. As this study was conducted in porcine epicardial adipose tissue, many of the genes have not been identified by homologous name and function to human genes and, as such, are shown by their Ensembl Stable Id. Refer to supplementary dataset 5.
