## Supplemental Methods for "Single-nucleus transcriptomics of epicardial adipose tissue from females reveals exercise control of innate and adaptive immune cells"

**Animal Experiments**

*Application of the experimental ameroid constrictor*

Animals were pre-anesthetized by intramuscular administration of glycopyrrolate (anticholinergic; 0.004 mg·kg^-1^), midazolam (benzodiazepine; 0.5 mg·kg^-1^) and ketamine (dissociative anesthetic; 20 mg·kg^-1^). Pigs were fitted with anesthesia masks and induced with 3% isoflurane. Animals were then intubated, and anesthesia was maintained with 2-3% isoflurane and auxiliary O_2_ during aseptic surgery. Immediately prior to surgery and during surgical recovery, animals received buprenorphine hydrochloride (analgesic; 0.1 mg·kg^-1^, i.v.) every 3-6 hr, as needed for pain relief. Antibiotic (ceftiofur 5 mg·kg^-1^, i.m. Excede®) was administered intramuscularly immediately prior to surgery. Furthermore, bupivacaine (local anesthetic) was applied intramuscularly to the third, fourth, and fifth intercostal spaces prior to cutting and topically along the suture line (2 cc per space) during closure. Pancuronium (neuromuscular blocker; 0.1 mg·kg^-1^) was administered intravenously immediately prior to opening the chest cavity to allow for placement of Finochietto retractors. During surgery, animals were given the following drugs as necessary: lidocaine (anti-arrhythmic; 1 mg·kg^-1^), epinephrine (adrenergic agent; 10 µg·kg^-1^), and atropine (antimuscarinic; 0.5 mg·kg^-1^). Upon opening of the pericardium, the proximal portion of the left circumflex artery was evaluated and an ameroid constrictor (2.5-3.5 mm) was placed to ensure a tight yet non-constrictive fit. The pericardium was closed and the thoracotomy was repaired in tissue layers. Immediately following surgery, Pigs received ketofen (non-steroidal anti-inflammatory; 3.0 mg·kg^-1^, i.v.). The ameroid constrictor progressively narrows the artery and induces an inflammatory response resulting in gradual occlusion, with total obstruction occurring at approximately two-three weeks post-operatively(1). Pigs recovered for eight weeks before the sedentary or exercise training experimental regimen was initiated.

*Sedentary and exercise protocols*

Speed and duration of the exercise training sessions were progressively increased so that during the last three weeks of training, animals ran at 4-5.5 mph for 60 minutes and at 6 mph for 5-15 minutes. Grade of the treadmill was maintained at 0% throughout the exercise bouts. The progressive nature of the exercise regimen was dependent upon the tolerance of each pig and therefore, ranges of speed and duration presented represent differing abilities of the animals. Exercise-trained and sedentary swine were fed once daily immediately after the exercise training session to serve as positive reinforcement for the exercise bout, with water provided ad libitum. Sedentary pigs maintained normal activity in their pens throughout the experimental protocol. Effectiveness of the exercise training regimen was determined by comparing heart-to-body weight ratio and skeletal muscle citrate synthase activity. Animal heart weight, body weight, heart-to-body weight ratio, and skeletal muscle citrate synthase values were compared using Student’s t-test. Data are presented as mean ± SEM, and *n* values in parentheses reflect the number of animals studied.

*Euthanasia for tissue collection*

Pigs were anesthetized with intramuscular administration of xylazine (2.25 mg·kg^-1^) and ketamine (35 mg·kg^-1^), fitted with an anesthesia mask, induced with 3% isoflurane and then intubated and a surgical plane of anesthesia was maintained with 3% isoflurane and auxiliary O_2_. A left lateral thoracotomy was performed in the fourth intercostal space, followed by administration of heparin (500 U·kg^-1^, i.v.). Hearts were removed, placed in Krebs bicarbonate buffer (0-4 °C) and weighed. Krebs bicarbonate buffer contained (in mM): 131.5 NaCl, 5 KCl, 1.2 NaH2PO4, 1.2 MgCl2, 2.5 CaCl2, 11.2 glucose, 13.5 NaHCO3 and 0.025 EDTA.

**Single-nucleus RNA sequencing**

For isolation of nuclei from adipose tissue, ~ 0.1 grams of snap frozen EAT was used for each nuclei isolation preparation. Samples were kept frozen and minced on a frozen cold block -20° C until the pieces were no larger than a grain of rice. The minced tissue was added to a 1.5 mL Eppendorf tube along with ice-cold Nuclei EZ Lysis Buffer (MilliporeSigma Cat # EZ Prep Nuc-101) supplemented with 0.2 – 0.5 U/uL RNAse inhibitor (Protector RNA Inhibitor, Millipore Sigma). Tissue was further homogenized in Lysis buffer using a pestel for 1.5 mL microcentrifuge tubes (USA Scientific 14155390). After mixing with a wide-bore pipette, the sample was allowed to incubate on ice for 5 minutes. After incubation the homogenate was passed through a 70µm strainer into a new tube and centrifuged at 500 x g at 4° C for 5 minutes. The remaining pellet was retained while the supernatant was discarded. The pellet was resuspended in G30 buffer (OptiPrep) using a wide-bore pipette tip and then the solution was underlaid with fresh G30 solution to establish a concentration gradient between the two. The tube was then centrifuged at 8,000 x g at 4°C for 20 minutes. After centrifugation, a nuclei pellet forms at the bottom of the tube while contaminating cell membranes and oil/fat remain in the suspension. Samples were washed and resuspended in ice-cold Nuclei Wash and resuspension buffer. Nuclei concentration/quality was assessed using the MoxiGo II Cell QC analyzer (Orflo, technologies). Nuclei were also assessed microscopically via trypan blue staining to visualize quality and the amount of residual debris within the sample.

Single nuclei 3` sequencing libraries were generated following the 10x Genomics dual index manual preparation v3.1 reagent kit with a target nuclei recovery of approximately 5,000 – 10,000 nuclei for each reaction. GEM partitioning was carried out in the 10x Genomics Chromium X system and following library construction following the manufacturer’s protocol. Sequencing libraries were quantified with the Qubit 4.0 Fluorometer high sensitivity dsDNA detection kit (Thermofisher). Library sizes were assessed using the Agilent TapeStation 4200, D1000 DNA tape system (Agilent Technologies). All sequencing library concentrations were normalized to 4 nM and pooled at equimolar ratios for sequencing on an Illumina NovaSeq 6000 2x150 sequencing run to generate a minimum of 200 million, paired-end sequencing reads for each sample. Sequencing reads were loaded into the 10x Genomics Cell Ranger software for sequence alignment, filtering, barcode and UMI counting.

**Bioinformatic analysis**

28,473 cells were profiled across 4 libraries from 4 tissue samples (tissues pooling from 2-3 swines), capturing 7,118 cells per library, 10,439 genes per cell, and 27,296 reads per cell on average. Low-quality cells and under expressed genes were excluded based upon four Quality control (QC) metrics: (i) genes with non-zero expression in fewer than 10 cells were excluded; (ii) cells with total UMI counts fewer than 200 were excluded; (iii) cells with percentage of reads mapping to mitochondrial genes more than 10% were excluded.

All 4 samples were integrated together to generate a merged Seurat Object, without any batch correction using Seurat v3 R Package. The data was Log-normalized using the “Simple Norm” method using a scale factor of 10,000. Principal component analysis (PCA) was performed to obtain the first 50 Principal components (PCs), of which the first 10 PCs were used to build a community of cells using k nearest neighbors (k=20) algorithm. The cells were further clustered using shared nearest neighbor information and modularity optimization using the Louvain algorithm as implemented in Seurat R Package v3. The preprocessed single cell data was visualized using Uniform Manifold Approximation and Projection (UMAP), a non-linear dimensional reduction technique. The differentially expressed marker genes for each cluster were identified using the Wilcoxon Rank Sum test. The obtained clusters were annotated based on their marker genes and cell types were identified using pig atlas (<https://dreamapp.biomed.au.dk/pigatlas/>). The scType database was used to further cluster and annotate the mesenchymal cells (2). An unknown cluster of cells was removed from the study as no significant markers corresponding to the atlas were found resulting in 20655 genes across 24382 cells for further analysis.

For within-and cross-tissue communication prediction, UMI count matrices and cell type/state assignment were exported for each cell as two input files for CellChat R package. CellChat is a publicly available repository of curated receptors, ligands and their interactions. CellChat R Package was used to infer cell-cell communication within and across cell types using the CellChatDB database. CellChatDB is a publicly available repository of curated receptors, ligands and their interactions. For each cell type, significantly differentially overexpressed ligand-receptor pairs were identified. Next, the communication likelihood was computed using the average expression values of a ligand in one cell type and that of a receptor/cofactor in another cell type. Significant interactions were pruned by permutation tests. An intercellular communication network was generated along with communication probabilities that measure the strength of relationships between each ligand-receptor pair. The communication network for each signaling pathway was generated by summing up all corresponding ligand-receptor interactions for that pathway. The CellChatDB was subset to specifically identify paracrine, endocrine and juxtacrine signaling pathways.
